# *Cryptococcus* displays spore-specific invasion of alveolar epithelial cells

**DOI:** 10.1101/2023.12.14.571430

**Authors:** Sébastien C. Ortiz, Rachael Fortune-Grant, Andrew J. Thom, Joshua Davies, Robin C. May, Rebecca A. Drummond, Margherita Bertuzzi

## Abstract

Human fungal pathogens, including *Cryptococcus neoformans,* cause 1.5 million annual deaths. *Cryptococcus* causes disease when it disseminates out of the lung and into the brain which can occur years after initial exposure (latency) via mechanisms that remain unknown. Spores of *Cryptococcus* display distinct surface epitopes, host-interactions, and disease kinetics to the vegetatively growing yeast morphotype, yet they remain understudied likely contributing to our lack of understanding of pathogenesis. One of the first barriers spores encounter are non-professional phagocytic <u>A</u>irway <u>E</u>pithelial <u>C</u>ells (AECs). Here, we demonstrated that *Cryptococcus* spores preferentially invade AECs both *in vitro* and *in vivo*. Once inside spores can germinate, subsequently replicate, persist and/or escape. This ability to enter AECs correlates with a preferential ability of spore to cross AEC barriers. Together our work indicates that AECs can be invaded by *Cryptococcus* spores and may serve as a previously ignored intracellular host niche, providing alternative hypothesis for both *Cryptococcus* dissemination and latency.

## Introduction

Invasive fungal diseases cause over 1.5 million deaths annually, with mortality rates often over 50% due to lack of diagnostics and antifungal treatments [1,2]. By understanding how fungi establish infections, persist in hosts and disseminate, we can develop much needed diagnostics, prevention, and antifungal therapeutics to combat invasive fungal disease. *Cryptococcus neoformans,* an inhaled facultative intracellular pathogen, is at the top of the WHO’s priority list of human fungal pathogens, largely due to the high mortality associated with cryptococcosis resulting in an estimated 181,000 annual deaths [3,4]. Key aspects of *Cryptococcus* disease progression are still not understood. Studies have focused on understanding the virulence of this pathogen and its ability to colonise the brain (the primary site of disease), but little is known about how *Cryptococcus* disseminates out of the lung (the initial site of infection). Additionally, via unknown mechanisms, *Cryptococcus* undergoes latency and is able to cause disease in hosts years or decades after initial infection [5,6]. Accordingly, after treatment for cryptococcosis, patients living with HIV/AIDS undergo a minimum of a year of secondary prophylaxis to prevent resurgence of the pathogen [7]. The mechanisms of *Cryptococcus* dissemination, latency and persistence remain unknown.

One of the potential reasons that the pathogenesis of *Cryptococcus* remains poorly understood is that the morphotype (morphologically distinct cell-type) used for the vast majority of studies is the yeast form; however, yeast are too large to reach the lower airway and thus unlikely to initiate infection under normal circumstance. *Cryptococcus* produces sexual spores (basidiospores), a dormant and stress resistant morphotype small enough to reach lower airways; consequently, spores are the presumed infectious morphotype in *Cryptococcus* disease [8,9]. Alas, difficulties associated with producing and working with *Cryptococcus* spores has resulted in a problematic knowledge gap regarding spore-mediated infections despite the fact that spores have distinct surface properties, interact with host immune cells differently than yeast, and, critically, disseminate out of the lung when yeast cannot [9–13]. The mechanisms of spore-specific dissemination out of the lung remain unknown, emphasising a need for the investigation of spore pathogenesis [13].

One of the first barriers inhaled pathogens encounter are non-professional phagocytic <u>a</u>irway <u>e</u>pithelial <u>c</u>ells (AECs). Alveolar AECs make up 24% of the lung parenchyma, likely have extensive and prolonged contact with inhaled fungal spores and are key for the antimicrobial defence of the lung, which if mis-regulated may lead to an opportunity for pathogens to persist [14–17]. Studies looking at the interactions of *Cryptococcus* with AECs show that yeast are not readily internalised [18–21]. Given the unique characteristics of spores and the importance of AECs in the host lung defence, we hypothesised that spores interact differently with AECs when compared to yeast and found that *Cryptococcus* spores invade AECs both *in vitro* and *in vivo*. Once inside, spores can germinate, replicate, persist and even escape. Furthermore, spores show preferential crossing of epithelial barriers, potentially implicating spore-AEC interactions in dissemination out of the lung. Our results identify AECs as a novel host cell-type that *Cryptococcus* can invade and exploit in a spore-specific manner. AECs also provide an intracellular environment which may facilitate latency/persistence of spores, a fundamentally dormant morphotype. These results provide viable alternative hypotheses to *Cryptococcus* dissemination, latency and persistence and emphasise the importance of evaluating different morphotypes in pathogenesis to open the door to novel opportunities for much needed diagnostic and therapeutic intervention.

## Results

### *Cryptococcus* spores, unlike yeast, are readily internalised by AECs and can germinate and replicate intracellularly

Previous studies investigating the ability of *Cryptococcus* yeast to enter AECs have shown that even at high infectious doses (Multiplicity of Infection (MOI) of 10) very few yeast cells are internalised (∼0%) [20]. Internalisation increases slightly for acapsular mutants, but these are avirulent and thus unlikely to be a cause of human infection [18–21]. Spores are known to have distinct surface properties that differ from those of yeast cells, suggesting that spores may interact with AECs differently from yeast [9–11, 22].

To test this, we exposed AECs to spores and carried out differential staining followed by imaging flow cytometry (IFC) (**Fig. 1**) [23]. A549 immortalised type-II alveolar epithelial cells were exposed to either mCherry (mCh) tagged *Cryptococcus* spores (produced from prolific sporulator strains JEC20xJEC21), yeast of the same background (JEC20+21), or yeast of the previously evaluated background (KN99α) at a low MOI of 0.167. Internalisation was evaluated 6 hours post infection, and counter staining with calcofluor white (CFW) was used to differentiate between AECs associated with *Cryptococcus* extracellularly versus AECs containing intracellular *Cryptococcus* (AEC**_i_**) (**Fig. 1A**). Consistent with previous studies, neither KN99α nor JEC20+21 yeast cells were taken up by AECs; however, JEC20×21 spores were readily internalised by AECs with 1.91% of AECs having 1 or more internalised spores (1.91% AEC**_i_**), and 20% of which had more than one spore per AEC (average stoichiometry of 1.34) (**Fig. 1B**) [20,21]. This rate of internalisation is equivalent to ∼15.4% of infectious inoculum being internalised by AECs within 6 hours.

**Figure 1.**
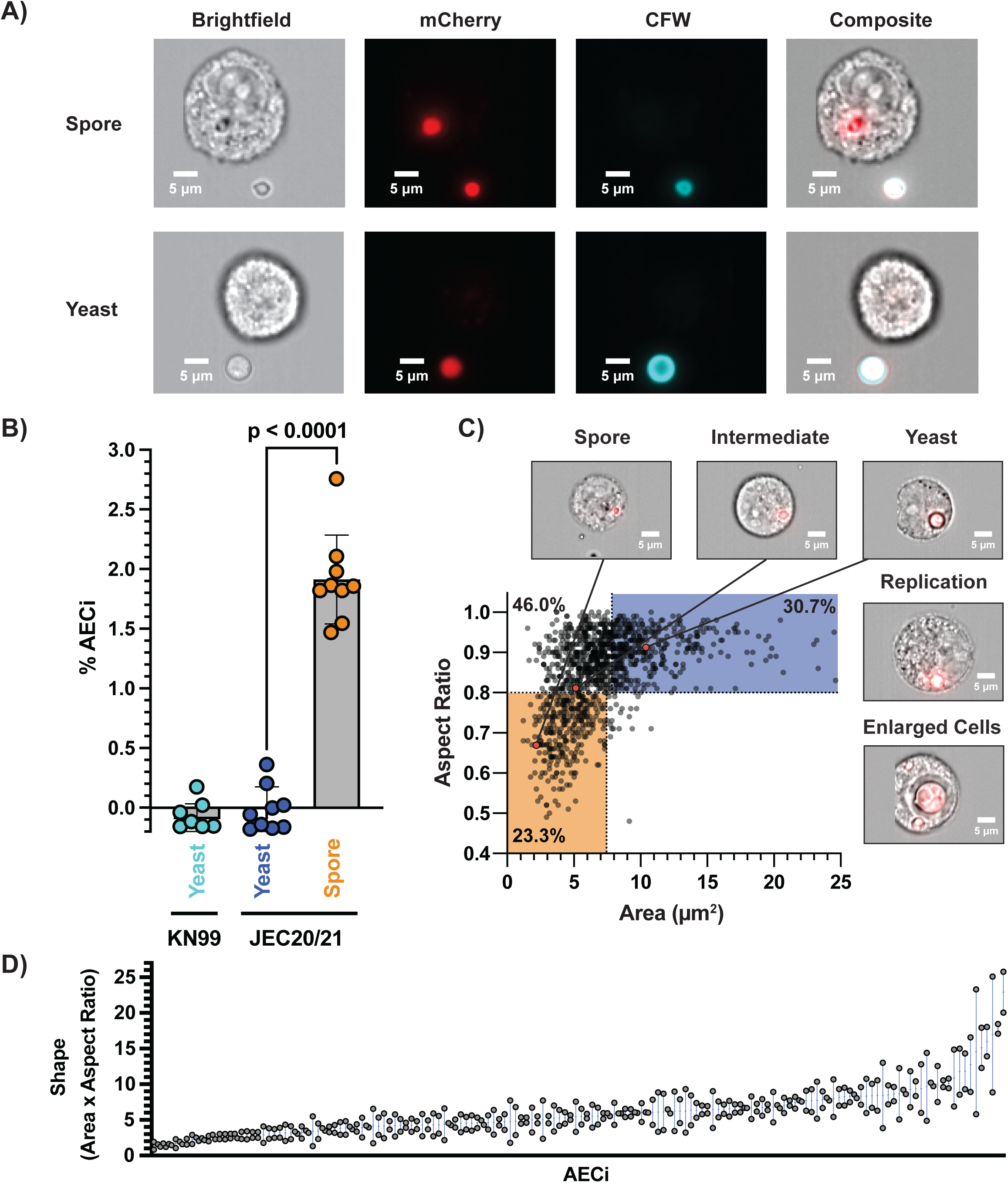
Spore-specific invasion of airway epithelial cells (AECs) by *Cryptococcus*. **A)** Representative images from Imaging Flow Cytometry (IFC), 6 hours post infection (hpi). **mCherry** tagged *Cryptococcus* used to identify AECs interacting fungal cells and Calcofluor White was used to distinguish internalised from non-internalised *Cryptococcus*. **B)** Percent of AECs with internalization events (AEC_i_) quantified for yeast (KN99 and JEC20/21), and spores (JEC20xJEC21), demonstrating that **spores** are readily internalised while **yeast** are not. A value of 1.91% AEC_i_ reflects 15.4% of spore infectious dose being internalised. One-way ANOVA p < 0.0001 **C)** IFC was performed 24 hours post infection to evaluate intracellular germination kinetics. Morphotypes were quantified based on cell size (surface area - µm^2^) and shape (aspect ratio), with spores being small and oval and yeast being large and round. Along with heterogeneous germination kinetics, both intracellular replication and cell enlargement were observed. **D)** Correlation of spore shape for *Cryptococcus* residing within the same AECs (Stoichiometry = 2) demonstrate a link between host AEC and germination kinetics p < 0.0001 two-way ANOVA.

To determine if the spore-specific internalisation observed was due to intrinsic spore properties rather than a generic induction of host-response to *Cryptococcus* spore challenge, internalisation was evaluated during a 6-hour co-infections with spores and yeast cells with different fluorophores (mCh and GFP) (**Supplementary Fig. 1A**). These experiments show that during spore/yeast co-infections, regardless of the fluorophore utilised, only spores can invade AECs. Spores tested are produced from congenic strain pairs and therefore have a homogeneous genotype; however, to ensure that the observed differences in internalisation were due to morphotype-specific differences in the properties of spores and yeast cells, rather than the result of a recombinant population, JEC20×21 spores were germinated into yeast (JEC20×21 yeast) and, upon infection of AECs, JEC20×21 yeast mirrored the lack of internalisation displayed by the JEC20+21 yeast populations (**Supplementary Fig. 1B**). These results support the idea that spore-specific invasion of AECs is due to intrinsic properties of spores rather than a generic induction of a host response or the recombinant nature of spores. The spore-specific ability to invade AECs was further validated in primary AECs (**Supplementary Fig. 1C**) and with serotype A KN99“a”xα spores (**Supplementary Fig. 1D**), confirming morphotype-specific invasion of AECs by spores but not the yeast counterpart.

In order to cause disease, spores need to germinate into vegetatively growing yeast. *Cryptococcus* spores readily germinate in the lung environment with cryptococci recovered from the bronchial alveolar lavage of mice 18 hours after infection with spores primarily being in the yeast form [13]. In the absence of AECs, spore populations were found to germinate fully by 24 hours in tissue culture conditions (37°C + 5% CO_2_) with a previously established bimodal phenotype (**Supplementary Fig. 2**) [24]. To determine if spores germinate within AECs, differential staining and IFC was performed on cells after 24 hours of infection with either spores or yeast (MOI of 1) and intracellular germination was evaluated by quantifying the size (surface area) and circularity (aspect ratio) of internalised *Cryptococcus* using previously establish metrics [25]. As observed previously, with this higher MOI, spores were readily internalised by AECs (12.6% AEC**_i_**) as opposed to yeast (0% AEC**_i_**) (**Supplementary Fig. 3Ai**). At 24 hours post infections, intracellular *Cryptococcus* was present as different morphotypes, i.e. 30.7% yeast, 46.0% intermediates, and a remaining 23.3% spores (**Fig. 1C**), indicating that spores readily germinate but that germination is inhibited in the intracellular environment. This population spread demonstrates a previously characterised asynchronous phenotype, indicating that internalised spores either react non-uniformly within the intracellular environment or that AECs have a range of intracellular environments that spores must germinate in [24]. These possibilities are not mutually exclusive. In addition to germination state, we observed that germinated spores can actively replicate 24 hours post-infection, as evidenced by daughter cell formation. Enlarged intracellular *Cryptococcus* yeast were also observed, consistent with the large size variation that is classically observed in lung environments [26]. The intracellular stoichiometry did not dictate the level of germination given that comparable germination states were observed across AEC**_i_** having internalised different numbers of spores (**Supplementary Fig. 4A**). On the other hand, when multiple spores were internalised by the same AEC**_i_**, intracellular spores displayed similar germination kinetics with similar shapes observed 24 hours post infection (**Fig. 1D**: Stoichiometry = 2, **Supplementary Fig. 4B**: Stoichiometry ≥ 3), suggesting that germination is linked to each internalisation event. Together these results show that spores can invade AECs, and once inside, can germinate heterogeneously, and subsequently replicate. This implicates AECs as a novel spore-specific intracellular niche that *Cryptococcus* can invade and proliferate within.

### Once germinated, *Cryptococcus* can undergo rapid replication, non-lytic escape, or persist inside AECs

To further examine the behaviour of internalised and germinated spores, fluorescence microscopy was used to monitor intracellular populations hourly from 6 to 26 hours post infection. CFW counterstaining was used at 6 hours post infection to demark internalisation events. We identified three behaviours that have previously been identified for intracellular yeast in macrophages, which are rapid replication (**Fig. 2A**), large vacuole formation (**Fig. 2B**) and non-lytic escape – also termed vomocytosis (**Fig. 2C**) [27–29]. Between 240 and 358 internalised *Cryptococcus* events per biological replicate were evaluated for the occurrence of budding, rapid replication balloon formation, vacuole formation, and non-lytic escape over 26 hours (**Fig. 2D)**. Intracellular budding occurred in 10.6% of internalisation events (21.0% of germinated spores), while rapid replication balloons occurred in 3.0% of events (6.3% of germinated spores) and was limited to when there was more than one fungal cell present within a single AEC. Vacuole formation occurred in 30.1% of germinated spores and only 2.0% of ungerminated spores (16.2% of all events). Finally, non-lytic escape was observed for 12.3% of germinated intracellular spores, and only 1.5% of non-germinated spores (7.0% of all events). These results demonstrate a large heterogeneity of intracellular behaviours, and that *Cryptococcus* can readily escape AECs once germinated.

**Figure 2.**
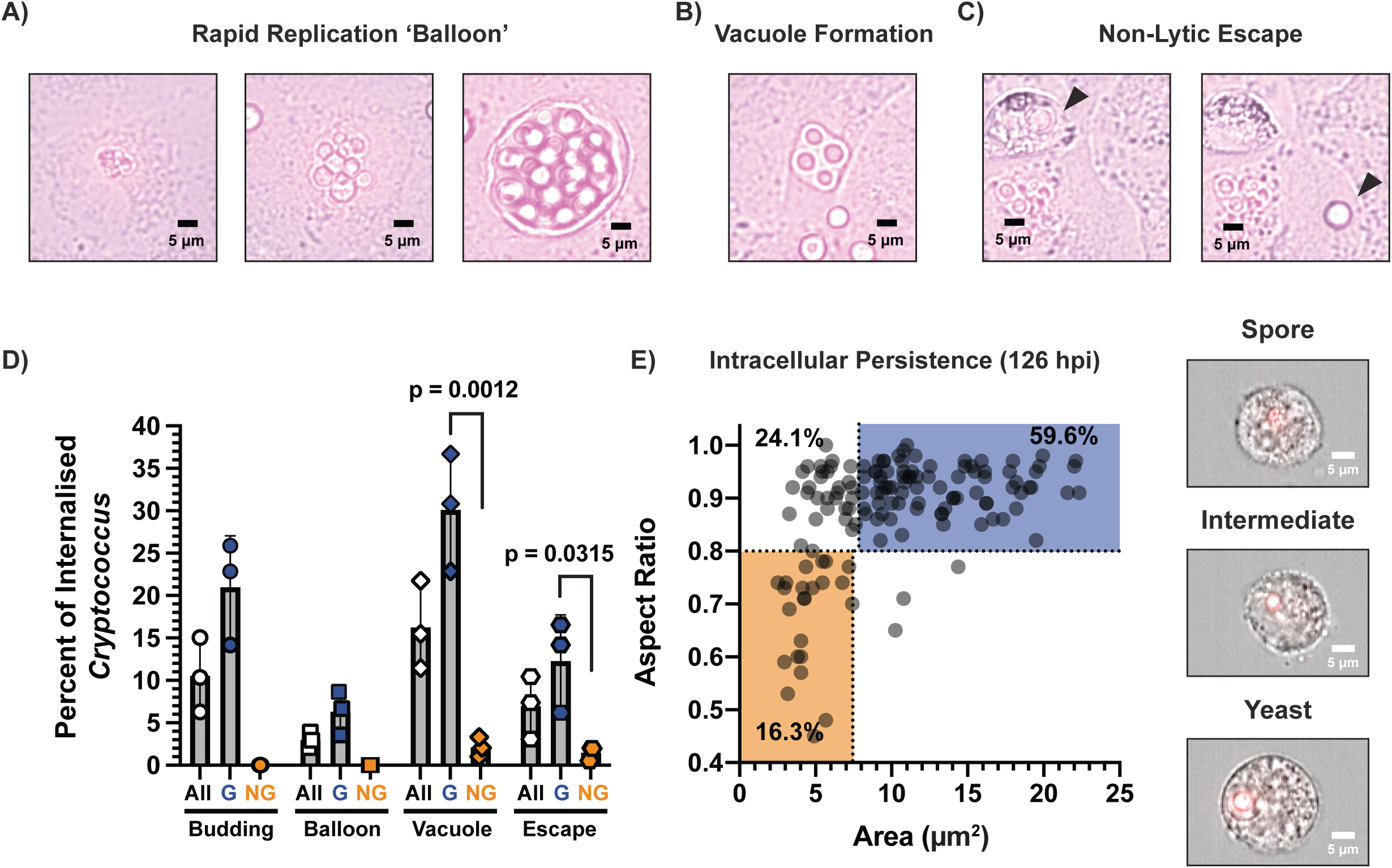
Internalised *Cryptococcus* spores display a variety of behaviours. Representative images of behaviours observed using time course microscopy of **A) Rapid Replication:** Internalised clusters of spores (6 hpi), efficiently germinating (16 hpi), and proceeded to rapidly replicate resulting in an AEC full of *Cryptococcus* yeast (24 hpi). **B) Vacuole formation** (25 hpi) **C) Non-lytic escape:** Internalised spore having germinated (24 hpi) undergoes non-lytic escape (25 hpi). **D) Quantification of intracellular behaviours:** Population percentages for All, Germinated (G) or Non-Germinated (NG) *Cryptococcus* that show the behaviours described: budding, rapid replication, vacuole formation, and non-lytic escape. Statistics from one-way ANOVA **E) Intracellular Persistence:** IFC identified intracellular *Cryptococcus* in spore infected cells 126 hpi displaying high levels of germination, and the presence of ungerminated spores and intermediates.

To investigate whether internalised *Cryptococcus* can persist within AECs for a prolonged period of time, A549 cells were infected with spores or yeast for 6 hours, then re-seeded to a low density to allow continue growth for an additional 5 days (120 hours). After 5 days, cells were evaluated for the presence of internal *Cryptococcus* cells. Once again, yeast-infected AECs had no intracellular *Cryptococcus* events; however, despite the daily duplication of AECs, we were still able to detect intracellular *Cryptococcus* in spore-infected AECs after 126 hours of infection (**Supplementary Fig. 3Aii**). The germination state of these internalised *Cryptococcus* cells was evaluated and a full range of germination morphotypes were identified with 59.6% yeast, 24.1% intermediates, and a remaining 16.3% spores (**Fig. 2E)**. While this population was more germinated than at 24 hours (up from 30.7% yeast), there were still spores and intermediates present after 126 hours. To determine the viability of intracellular *Cryptococcus* 126 hpi, internalisation events were sorted using Fluorescence assisted cell sorting (FACS) and grown in liquid YPD for 1 week. Viability was evaluated for 864 internalization events across three technical and three biological replicates (96/replicate) and the presence of fungal growth showed that 79.25% (StDev: 5.60) of AECs with internalisation events had viable *Cryptococcus* capable of growth under these conditions. It is formally possible that *Cryptococcus* escapes these AECs and re-enter; however, non-lytic escape was rare for non-germinated spores (1.5%) unlike germinated spores (12.3%) (**Fig. 2D)** and we have shown that once germinated into yeast, *Cryptococcus* lose its ability to invade AECs (**Supplementary Fig. 1B**). Additionally, AECs do not show significant signs of apoptosis or necrosis after 6, 24, and 126 hours of infection, likely due to the low virulence of JEC20/JEC21, making lysis and re-entering much less likely than intracellular persistence (**Supplementary Fig. 3B**) [30]. Together, these results demonstrate that AECs provide an intracellular environment in which spores can germinate, replicate, escape and even persist.

### *Cryptococcus* spores preferentially cross AEC barriers and are internalised *in vivo*

Spores of avirulent yeast preferentially disseminate out of the lung resulting in disease in intranasal murine models of infection [13]. Given the unique ability of spore to enter AECs, and subsequently germinate, replicate and escape, we hypothesised that spore-specific invasion of AECs would enable them to cross epithelial barriers better than yeast. To test this, we monitored the crossing of *Cryptococcus* across trans-wells seeded with Calu-3 bronchial-like cell line, which unlike A549 cells, form tight junctions and impermeable barriers. Spores are internalised by bronchial Calu-3 cells but less efficiently than alveolar A549 cells (**Supplementary Fig. 3Aiii**). Confluent Calu-3 cells were infected with either spores or yeast (MOI 5) and media from the lower compartment was plated for colony forming units at 0, 24 and 48 hours (**Fig. 3A**). After 48 hours, spores showed preferential crossing as compared to yeast, with 88.9% spore infected wells showing crossing, as opposed to 22.2% of yeast infected wells. The impermeability of Calu-3 transwell barriers was evaluated using a previously defined methodology which utilizes fluorescence monitoring of Dextran 70kDa-TexasRed crossing [**Supplementary Fig. 5A**] as a proxy of epithelial barrier integrity, along with transepithelial resistance measurements [**Supplementary Fig. 5B**] [31]. Provided that the addition of Dextran 70kDa-TexasRed did not alter the preferential *Cryptococcus* spore crossing observed [**Supplementary Fig. 5C**], these measurements confirmed the integrity of the transwell barrier beyond 48 hours post infection. Co-infections with mCh-tagged spores and GFP-tagged yeast also displayed preferential spore crossing, further supporting that spores are better able to cross an impermeable lung epithelial barrier than yeast [**Supplementary Fig. 5D**].

**Figure 3.**
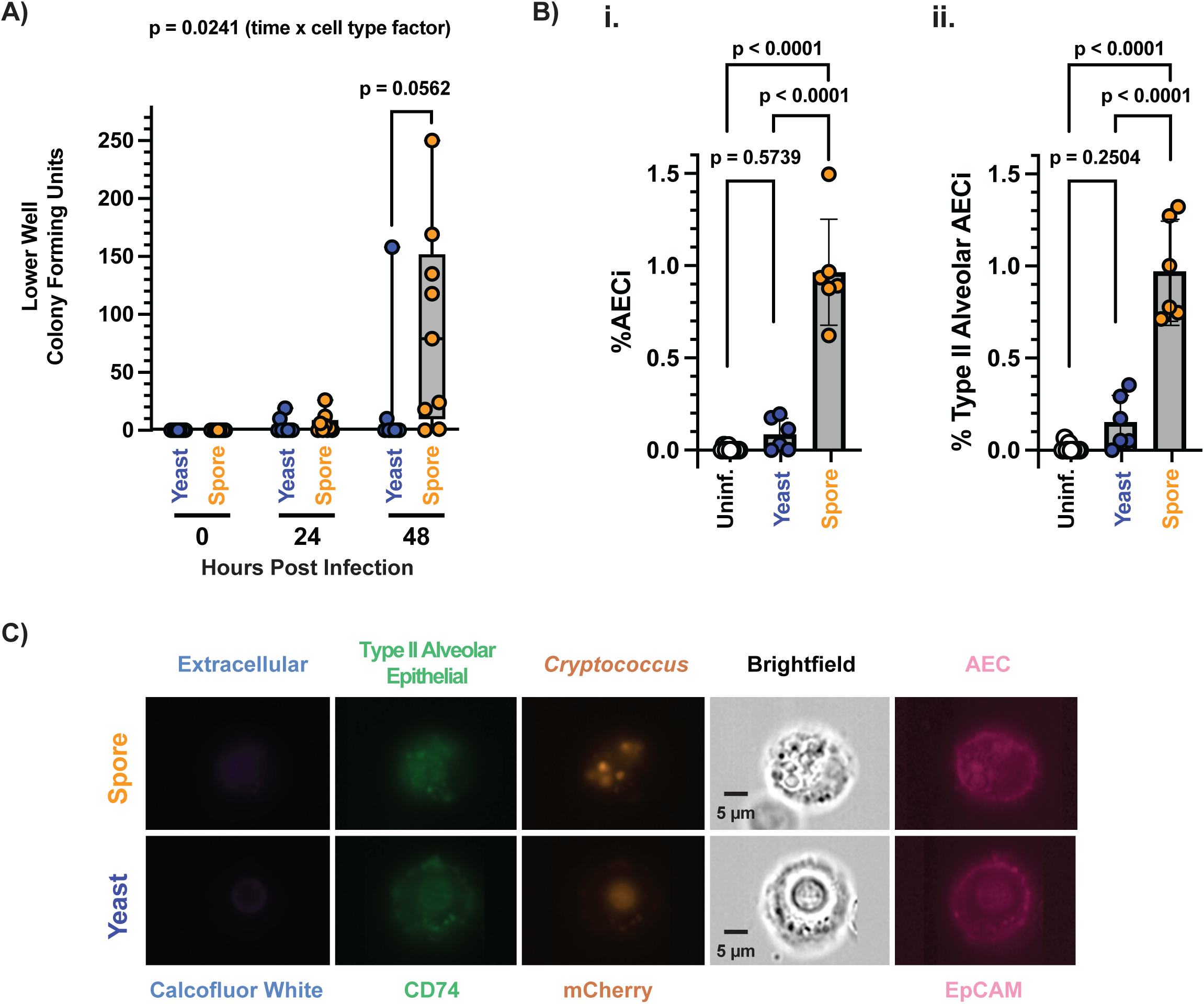
*Cryptococcus* spores preferentially cross AEC barriers and are internalised by AECs in a murine model of infection. **A)** Lower well colony forming units at 0, 24, and 48 hours in spore and yeast infected trans-wells indicating the crossing of AEC barrier. Statistics from 2-way ANOVA **B)** Percentage of i) AECs (EpCAM+) or ii) Type II alveolar epithelial cells (EpCAM+, CD74+) with intracellular *Cryptococcus* (mCherry+, CFW-), derived from mouse lungs 8 hours post intranasal infection with spores or yeast. All statistics derived from one-way ANOVA across samples. **C)** Representative Imaging Flow Cytometry panels of *Cryptococcus* cells (both Spores and Yeast) inside type II airway epithelial cell, 8 hours post intranasal infection. **Ch1: Calcofluor White** staining differentiates between fungal cells inside and outside AECs. **Ch2: CD74** staining is used to identify Type II alveolar AECs. **Ch4:** Nuclear localized **mCherry** tag identifies fungal cells. **Ch5:** Brightfield image of cells. **Ch6: EpCAM** staining is used to identify AECs.

Given this ability of spores to invade, persist inside, and escape AECs, we evaluated the *in vivo* internalisation of *Cryptococcus* by AECs using an intranasal murine model of cryptococcosis. Mice were intranasally infected with 5×10^6^ spores or yeast (JEC20mCh/JEC21mCh) and the infection was allowed to progress for 8 hours. This timepoint was chosen to ensure morphotype-specific characterisation because 8 hours is not long enough for spores to have germinated into yeast. Lungs were harvested, processed, pooled and differentially stained using our previously established methodology to identify AECs via IFC (**Fig. 3B&C**) [17]. We found that for spore-infected mice, 0.96% of recovered AECs (EpCAM^+^) and 0.97% of recovered type-II alveolar AECs (EpCAM^+^, CD74^+^) had internalised spores, with the majority of events having more than one spore (58.6%) with an average stoichiometry of 2.28 spores/AEC**_i_** (**Supplementary Fig. 4Ciii**). On the other hand, for yeast-infected mice, only 0.09% of recovered AECs and 0.15% of recovered type-II alveolar AECs had internalised yeast, with 21.3% having 2 yeast cells with an average stoichiometry of 1.14 yeast/AEC**_i_**. These data indicate that spores are able to cross epithelial barriers *in vitro* and invade AECs *in vivo* better than yeast. These results reveal AECs as a novel intracellular host cell type that *Cryptococcus* can invade and offers potential alternative mechanisms of spore-specific dissemination and persistence within the host lung.

## Discussion

*Cryptococcus* spores, along with ‘desiccated’ yeast, are the presumed infectious particles in cryptococcal disease; however, vegetatively growing yeast are not generally considered to be the primary infectious propagules due to their large size preventing them from reaching the lower airways. Contrary to this, and likely due to the difficulties associated with working with spores and the low viability of ‘desiccated’ yeast, the vast majority of studies on *Cryptococcus* host-pathogen interactions and disease kinetics have been performed with vegetatively growing yeast and not spores. While understanding yeast-host interactions is critical, ignoring other morphotypes prevents us from understanding the complete picture concerning *Cryptococcus* infections and subsequent disease. By neglecting to study these morphotypes, we miss out on opportunities to develop diagnostics, prevention and treatments for fungal disease. In this study, we demonstrate that spores, which have distinct surface properties, interactions with host cells, and disease kinetics to yeast, are internalised by AECs when yeast are not [9–13]. Once inside, these spores can germinate, replicate, escape and/or persist within AECs (**Fig. 4**). These spore-specific host-pathogen interactions may have major implications for our understanding of both dissemination and latency.

**Figure 4.**
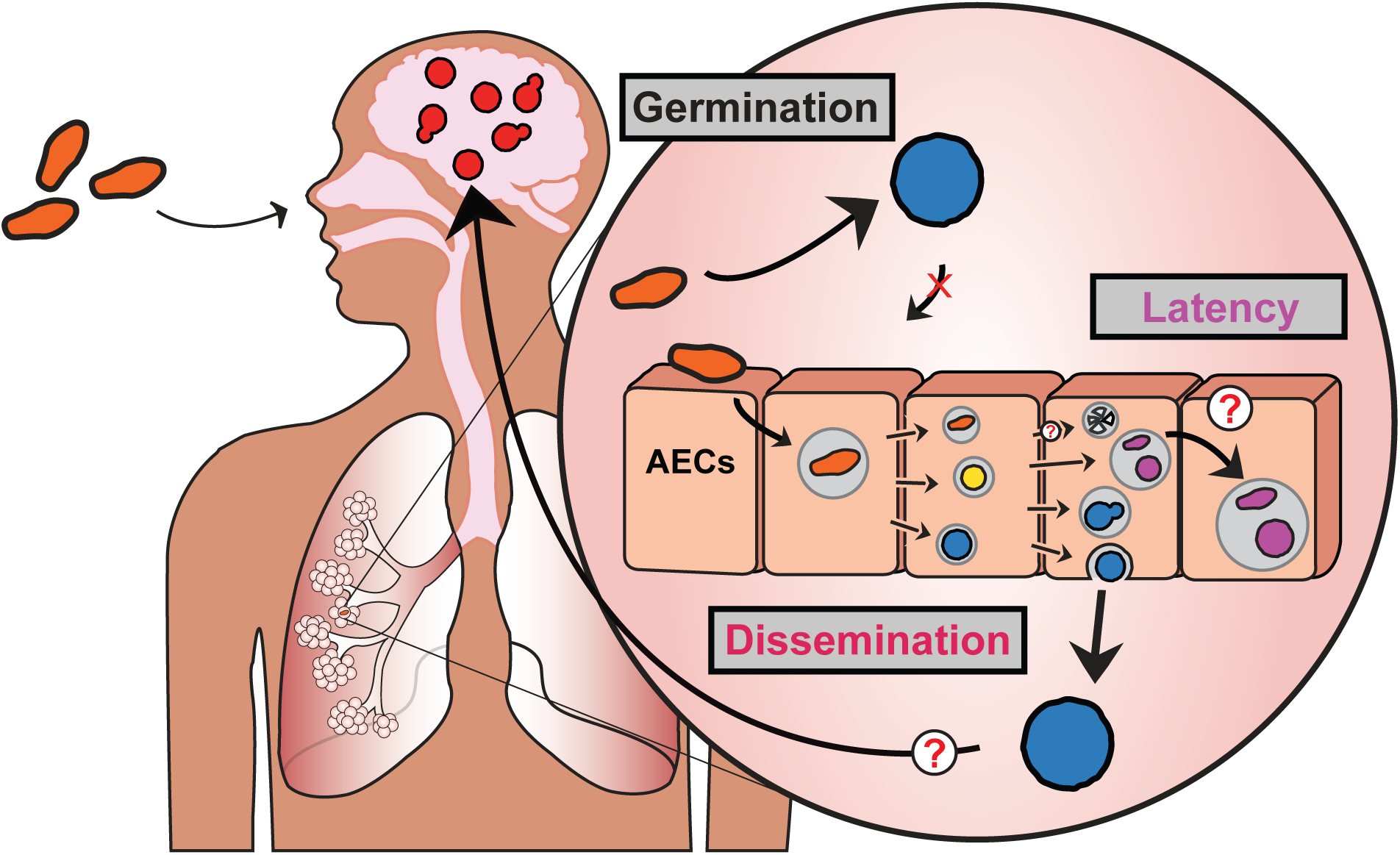
Schematics of our current understanding of *Cryptococcus*-AEC interactions. In this work we’ve demonstrated that spores, unlike yeast, are readily internalised by airway epithelial cells. This spore-specific interaction provides a unique intracellular niche where spores can germinate, and once germinated, can replicate, persist and non-lytically escape. The morphotype specific ability to enter AECs, survive, and escape provides a viable explanation for *Cryptococcus* spore-specific dissemination out of the lung. The ability of *Cryptococcus* to persist inside AECs, not only as fully germinated yeast, but as apparently dormant (ungerminated) spores provides an unexplored potential mechanism of latency.

Work by Walsh et al., demonstrated that spores of avirulent yeast (B3501/02) caused disease in an intranasal murine model 50 days after initial infection, while yeast did not [13]. Spore-specific virulence was directly linked to the ability of spores to disseminate out of the lungs when yeast could not, with dissemination to the kidneys within 6 days post infection, and eventually dissemination to the brain. Since spores readily germinate within 18 hours in the mouse lung, this work suggested that early host-pathogen interactions in the lung dictate spore-specific dissemination and disease. The authors postulated that spore-specific dissemination could result from a Trojan-Horse mechanism where spores hide inside macrophages to get shuttled out of the lung, a prominent hypothesis for yeast dissemination as well. The work demonstrated that spores were more efficient at initially reaching lung draining lymph nodes than yeast, and that trafficking to the lymph nodes was CD11c^+^ dependent. Unfortunately, the authors were unable to evaluate the role of macrophage uptake in further dissemination to other extra-pulmonary organs. The data we present here provides an alternative, but not mutually exclusive, hypothesis for the mechanism of dissemination out of the lung. Rather than a macrophage driven Trojan-Horse model, we propose spores may be able to disseminate out of the lung through transcytosis, where spores are internalised by epithelial cells, germinate, and non-lytically escape on the other side of the epithelial barrier. This alternative mechanism is further supported by recent work that identified the infrequent occurrence of ‘free floating’ yeast in the vasculature (large blood vessels) of mouse lungs 7 days post intranasal yeast infection, consistent with the low levels of internalisation by AECs we observed in our murine yeast infections [32]. These two mechanisms may even work in tandem, as reported with *Mycobacterium tuberculosis (Mtb)*, with monocytes containing *Mtb* more efficiently translocating across an epithelial barrier when AECs were also infected with *Mtb* [33]. In order to conclusively determine the precise role of both macrophages and AECs in *Cryptococcus* dissemination out of the lung, the molecular mechanisms driving spore-host interactions need to be identified to subsequently modulate spore-host interactions.

The first epidemiological evidence of *Cryptococcus* latency was obtained in 1999 [34]. Since this first study, a variety of evidence has supported the idea of latency/persistence in cryptococcal disease [5,6]. Relatively little is known about the mechanisms underlying latency/persistence; however, granulomas have largely been presumed to be the predominant source of latent *Cryptococcus* [35]. This idea originally stems from a comparison to *Mtb*, where similar granulomas have been the predominant paradigm for a mechanism of latency [5, 35, 36]. However, this has recently been challenged with evidence of *Mtb* DNA being present in non-professional phagocytic lung cells in the absence of tuberculous legions [36,37]. The frequency and importance of granulomas in Cryptococcal disease is unclear, and while granulomas are the presumed latency reservoir in the lung, there may be other mechanisms driving latency. Based on our data we propose an alternative mechanism where AECs could provide an intracellular environment, shielded from immune cells, in which *Cryptococcus* spores could lie dormant for extended periods of time. The presence of *Cryptococcus* cells within AECs of patients has yet to be investigated and these events would not be observed without actively looking for them.

Dormancy in *Cryptococcus* is poorly understood and the morphological state of *Cryptococcus* during latency is unknown. *In vivo* models have demonstrated the presence of low-metabolic activity yeast that match a dormant population [38]. Following this, *in vitro* conditions were developed to mimic this dormant population resulting in viable but non culturable cells (VBNC) [39]. Further research is required to properly understand the importance and role of this yeast subpopulation in *Cryptococcus* disease kinetics [6,38,39]. Importantly, spores are the classical example of a dormant particle, and for *Cryptococcus* these are exclusively produced in the environment, not inside the host. Spores shows distinct nutrient utilisation to yeast but the molecular mechanisms that control the escape from dormancy still require characterisation [8,11]. It is possible that spores, being a fundamentally dormant morphotype, are more readily able to remain dormant (or re-establish dormancy) in a host than yeast. The identification of AECs as a spore-specific intracellular environment, where spores can persist and germinate to varying degrees, provides a novel alternative for understanding latency and persistence in cryptococcal disease.

A growing body of evidence shows an important role for AECs in the antimicrobial potency of the lung, where the interactions of these non-professional phagocytes with pathogens can be either a benefit or detriment to the host [15,16]. This likely depends on both pathogen factors and host predispositions. The role of AECs in cryptococcal disease has largely been ignored which may be due to the lack of internalisation of the yeast morphotype. It will be interesting to understand if other understudied morphotypes present in the lung, such as VBNC yeast, seed cells, and Titanides, could interact similarly with AEC as spores do. This study expands on the already long list of differences between *Cryptococcus* spores and yeast and points towards AECs as a novel intracellular environment for the field to consider when studying *Cryptococcus* disease kinetics and potential therapeutics.

## Methods

### Fungal Strains and Strain Manipulation

*Cryptococcus neoformans* serotype D (*deneoformans*) strains JEC20-mCh (CHY4031), JEC21-mCh (CHY4028), JEC20-GFP (CHY3955), JEC21-GFP (CHY3952), and *Cryptococcus neoformans* serotype A strain KN99α-mCh were handled using standard techniques and media as described previously [13,40,41]. *Cryptococcus* spores were isolated from cultures as described previously [9]. Briefly, yeast of both mating types (JEC20-mCh and JEC21-mCh) were grown at 30°C for 2 days on yeast-peptone-dextrose (YPD) agar, resuspended in 1X phosphate buffered saline (PBS) [Sigma-Aldrich: D8537], mixed in a ratio of 1:1, and intermittently spotted onto dried V8 pH 7.0 agar plates. Plates were incubated for 7 days at 25°C in the dark and crosses were resuspended in 75% Percoll® [Sigma-Aldrich: GE17-0891-01] in 1X PBS for separation via gradient centrifugation (2110xg, 25 min). Spores were recovered, counted using a hemocytometer and assessed for purity by visual inspection. Germinated spores (JEC20×21 yeast) were generated by growing purified spores at 30°C for 3 days on YPD agar.

To produce mCherry tagged KN99“a”xα spores, a KN99“a”-mCh strain was derived by mating KN99α-mCh yeast with KN99“a” yeast on Murashige and Skoog pH 5.8 agar plates for two weeks. Spores were grown out for single colonies on YPD, screened for fluorescence to confirm mCh signal, and evaluated for mating ability with KN99α to identify “a” mating type. For infections KN99“a”xα spores were derived as described above, but a 90% Percoll® was used for gradient centrifugation, and due to 65% spore:yeast purity, inoculum was adjusted to account for yeast presence.

### Epithelial Cell Culture

The cell lines used in this study were A549, human pulmonary carcinoma epithelial cells [ATCC: CCL185] and Calu-3, human pulmonary adenocarcinoma epithelial cells [ATCC: HTB-55]. A549 cells were maintained in DMEM [Sigma-Aldrich: D5796] supplemented with 10% Gibco^TM^ FBS [ThermoFisher Scientific: A5256801] and 1% penicillin/streptomycin cocktail [Sigma-Aldrich: P0781] (DMEMs). Calu-3 cells were maintained in DMEM/F12 [ThermoFisher Scientific: 11320033] supplemented with 10%FBS, 1% penicillin/streptomycin cocktail (DMEM/F12s). Primary human AECs were purchased from Lonza (Healthy = 2547, TAN 43140 and 40273) following isolation from the distal portion of the human respiratory tract in the 1 mm bronchiole area. Primary AECs were maintained in Small Airway Epithelial Cell Growth Medium (Promocell) and used for infections within the ninth passage to avoid senescence. All cells were grown and maintained at 37°C, 5% CO_2_, with all passaging and seeding performed with Trypsin-EDTA [Sigma-Aldrich: T3924] mediated detachment.

### Flow Cytometry

#### Differential Fluorescence Imaging Flow Cytometry

Differential fluorescence IFC was performed following published methodology [23]. Briefly, confluent monolayers of A549 (or primary cells) cells across triplicate wells of 6-well plates [Greiner: 657160] were infected with either spores (JEC20-mChxJEC21-mCh, KN99“a”-mChxKN99α-mCh), yeast (JEC20-mCh+JEC21-mCh, KN99α-mCh, KN99“a”+KN99α) or germinated spores (JEC20×21 yeast) at varying MOIs (0.167, 1 or 5) for different lengths of times (6 hours, 24 hours). All experiments had uninfected controls processed identically as negative controls and for gating purposes. For yeast/spore co-infections A549 cells were infected at an MOI of 0.167 (per morphotype) with either spores (JEC20-mChxJEC21-mCh or JEC20-GFPxJEC21-GFP), yeast (JEC20-mCh+JEC21-mCh or JEC20-GFP+JEC21-GFP) or combinations of 2 fluorescently distinct cell types. For intracellular persistence evaluation, after 6 hours of incubation A549 cells were reseeded to attain confluency 120 hours later (effectively 126 hpi). At the end of the infections, A549 cells were trypsin digested and collected via centrifugation at 500x g. Cells were incubated in 500 µL of Binding Buffer, 1X (5 µL) Annexin V-FITC [Abcam: ab14085], 100 nM TO-PRO^TM^-3 [ThermoFisher Scientific: T3605] and 8 µg/mL Calcofluor White (CFW) [Sigma-Aldrich: F3543] for 5 minutes, centrifuged, and washed with 1X PBS. Cells were resuspended and evaluated on a Amnis ImageStream MkII. For each sample, 5000 single cells in focus were analysed across three independent acquisitions in biological triplicate. Analysis was performed for the identification of internalisation and host cell death using IDEAS® software according to previously defined analysis pipeline [24]. Uninfected A549 samples were used for background subtraction, and all samples were normalised based on infectious inoculum as determined by triplicate CFU counts. Visualisation of internalisation events was performed on both IDEAS® software and Fiji/ImageJ to quantify stoichiometry and germination. *Cryptococcus* germination quantification was performed by evaluation of area and aspect ratio of internalised *Cryptococcus* cells following previous established cut offs (Spores: Area <7.44 µm^2^, Aspect Ratio <0.8 | Yeast: Area >7.84 µm^2^, Aspect Ratio >0.8) as defined by Barkal et al. 2016 [25] across all biological replicates with technical replicates acquired for internalisation evaluation.

#### Fluorescence assisted cell sorting (FACS)

FACS (using a BD Influx Cell Sorter) was used to sort persistent *Cryptococcus* 126 hours post infection with spores. Infections and processing were performed as detailed above and single A549 cells containing intracellular *Cryptococcus* (mCh^+^/CFW^−^) were sorted into wells of 96-well plates containing YPD for 3 technical replicates for each of 3 infections (biological replicates). Plates were incubated at 30°C for 1 week and wells were visualized under a microscope and scored for presence of *Cryptococcus* growth.

### Microscopy

#### Intracellular Cryptococcus Behaviours

A549 cells were seeded into 24-well glass bottom plates [Greiner: 662892] for two days until reaching confluency, infected with JEC20-mChxJEC21-mCh spores at an MOI of 1. After 6 hours, media was removed and cells were stained with 8 µg/mL CFW for 5 minutes, washed with 500 µL 1X PBS and resuspended in Gibco^TM^ FluoroBrite^TM^ DMEM media [ThermoFisher Scientific: A1896701] supplemented with 10% FBS and 1% penicillin/streptomycin cocktail. All images were acquired on a Nikon Eclipse Ti Inverted Microscope, using a Nikon CFI Plan APO λ 40X objective and Hamamatsu ORCA-Flash4.0 LT C11440 camera, at 37°C and 5% CO_2_ for 20 hours after staining. Images were acquired across 3 biological replicates, 3 wells per replicate (10 positions/well) in bright field, 400 nm (100 ms exposure) and 550 nm (300 ms exposure). For each biological replicate, 240-348 internalisation events (determined by CFW-signal at 6 hours post infection and localisation) were monitored hourly from 6 to 26 hours post infection. For each internalised *Cryptococcus* event, germination (determined by size/shape), replication (determined by the addition of intracellular *Cryptococcus* cells i.e. budding), rapid replication (determined by number of intracellular *Cryptococcus* cells at least tripling by 26 hours and visual ballooning), vacuole formation (determined by large ring surrounding intracellular *Cryptococcus* cells), and non-lytic escape (determined by intracellular *Cryptococcus* cells floating away from host cells) were evaluated and quantified.

#### Quantitative Germination Assay

Germination of spores was evaluated across 24 hours in tissue culture conditions (supplemented DMEM, 37C, 5%CO_2_), and images were acquired on a Nikon Eclipse Ti Inverted Microscope, using a Nikon CFI Plan Fluor ELWD 20X objective and Hamamatsu ORCA-Flash4.0 LT C11440 camera. Germination was quantified using the previously published quantitative germination assay and custom algorithms [24,25]. Briefly, image analysis was conducted using a previously published custom FIJI macro to outline fungal cells which provide measurements of size and shape. The resulting data is subsequently organized using a custom Matlab script and categorizes cells with an area < 7.44 µm^2^ and aspect ratios of < 0.8 as spores, cells with an area > 7.84 µm^2^ and aspect ratios of > 0.8 as yeast and anything in between as intermediates. Data is plotted as 2-dimentional histograms of size and shapes to visualize germinating population data.

### Trans-well Crossing

Trans-well crossing experiments were performed by seeding Falcon® permeable support for 12-well plate with 8.0 µm transparent PET membrane [Corning: 353182] and associated companion plate [Corning: 353503] with Calu-3 cells at 1×10^6^ cells/well. Cells were allowed to reach confluency (8-12 days) as determined by a resistance measurement of >900 ohm using Milipore Millicell® ERS-2 Epithelial Volt-Ohm Meter and Milicell® ERS-22 Adjustable electrode set [MERSSTEX03]. Each upper well of the trans-well was infected with either spores or yeast (MOI of 5) and crossing was evaluated by plating lower well volumes onto YPD plates at 0, 24 and 48 hours post infection. YPD plates were incubated for 3 days, and total number of colonies/well/day were quantified. Impermeability of barrier was further confirmed by lack of fungal crossing following infection and resistance was monitored across all time points. Crossing was evaluated for 9 replicates of both spore and yeast infected wells.

In addition to the transepithelial resistance measurements used to evaluate impermeability, supplemental experiments as from previously published methodology [31] were performed using 70kDa Dextran-Texas Red [ThermoFisher: D1830], which was added to the upper wells at a concentration of 0.1 µM. Dextran crossing was monitored by evaluating the fluorescence (595/615 nm) of the lower well media every 24 hours post infection. Fungal crossing experiments were repeated, using the methodology above, in the presence of Dextran with spores (JEC20×21-mCh), yeast (JEC20+21-GFP) and with co-infections of both. Colonies from co-infection experiments were evaluated for either mCh or GFP signal to determine if they were spore-derived or yeast-derived *Cryptococcus* crossing event, respectively.

### Murine infection, lung processing, and staining for imaging flow cytometry

Murine infections were performed under UK Home Office Project License PBE275C33 at the Biomedical Service Unit, University of Birmingham. Mice were housed in individually ventilated cages under specific-pathogen free conditions, with access to standard chow and drinking water *ad libitum* under 12-hour light/dark cycle at 20-24°C and 45-65% humidity. Spores (JEC20-mChxJEC21-mCh) and yeast (JEC20-mCh+JEC21-mCh 50:50 mix) were enumerated using an haemocytometer and diluted in 1X PBS. Three groups of 8-10 week-old female C57BL/6NCrl mice (Charles River) were left uninfected (6 mice), or intranasally infected with spores (4 mice), or yeast (4 mice) at a final dose of 5×10^6^ cell/mouse. Mice were humanely euthanised by cervical dislocation at 8 hours post infection and lungs were perfused with 2 mL of 0.1 mg/ml Dispase [Sigma-Aldrich: D4693] in HANKS buffer [Sigma-Aldrich: H9269] directly into trachea. Lungs were processed as previously described [18]. Tissue digestion of whole lungs was carried out at 37°C for 1 hour with gentle rotation (100 rpm). Digestion buffer (0.1 mg/ml Dispase, 0.4 U/ml Liberase TL [Sigma-Aldrich: 5401020001] and 160 U/ml DNase [Sigma-Aldrich: D5025] was added and lungs were incubated an additional 10 minutes. Samples were diluted with 1X PBS + 2% FBS + 2 mM EDTA, systematically filtered through 100 µm, 70 µm and 40 µm filters [Sigma-Aldrich: 431752, 431751, 431750] and centrifuged at 500x g for 6 minutes at 4°C. Supernatant was removed, and pellet was resuspended in 10 mL of 1x RBC lysis buffer [BioLegend: 422401], incubated at 25°C for 5 minutes. Samples were pooled together (2 mice per pool), diluted with 1X PBS + 2% FBS + 2 mM EDTA, centrifuged for 6 minutes at 500x g and 4°C, supernatants were discarded, and the pellets were resuspended in remaining volume (∼200 µL). Samples containing excess remaining red blood cells post lysis were not evaluated via IFC. Each sample was incubated with 5 µg/mL anti-mouse CD16/32 [BioLegend, 101302] for 10 minutes at 4°C, centrifuged at 500x g for 6 minutes at 4°C and pellets were resuspended in 100 µL of 1X PBS + 2% FBS + 2 mM EDTA. Samples were incubated with 1 µg/mL Ep-CAM (CD326) PE/Cy7 Antibody [BioLegend, 118215] and 1 µg/mL CD74 FITC Antibody [Santa Cruz Biotechnology, sc-6262-FITC] for 25 minutes at 4°C, followed by 8 µg/mL CFW for 5 minutes. Samples were centrifuged at 500x g, for 3 minutes and resuspended in 100 µL of 1X PBS + 2% FBS + 2 mM EDTA prior to evaluation on an Amnis ImageStream MkII. For each pooled sample (three uninfected, two yeast infected, and two spore infected), 3 acquisition of 3000-4000 EpCam^+^ single cells in focus were taken as technical replicates. Data analysis was performed using the IDEAS® software and the pipeline previously described, using single stain controls for gating purposes.

### Statistical analysis of data

GraphPad Prism was used to interpret data and p values were calculated through unpaired t tests or, ordinary 1-way or 2-way ANOVA as specified in the figure legends. Error bars show standard deviation.

## Acknowledgments

We would like to thank Dr Christina Hull for the gift of the JEC20-mCh, JEC21-mCh, JEC20-GFP, and JEC21-GFP strains, Dr Gareth Howell and his team at the Flow Cytometry Core Facility, University of Manchester, and Ferdus Sheik at the Flow Cytometry Facility, University of Birmingham for technical support.

---

**Supplemental Figure 1.**
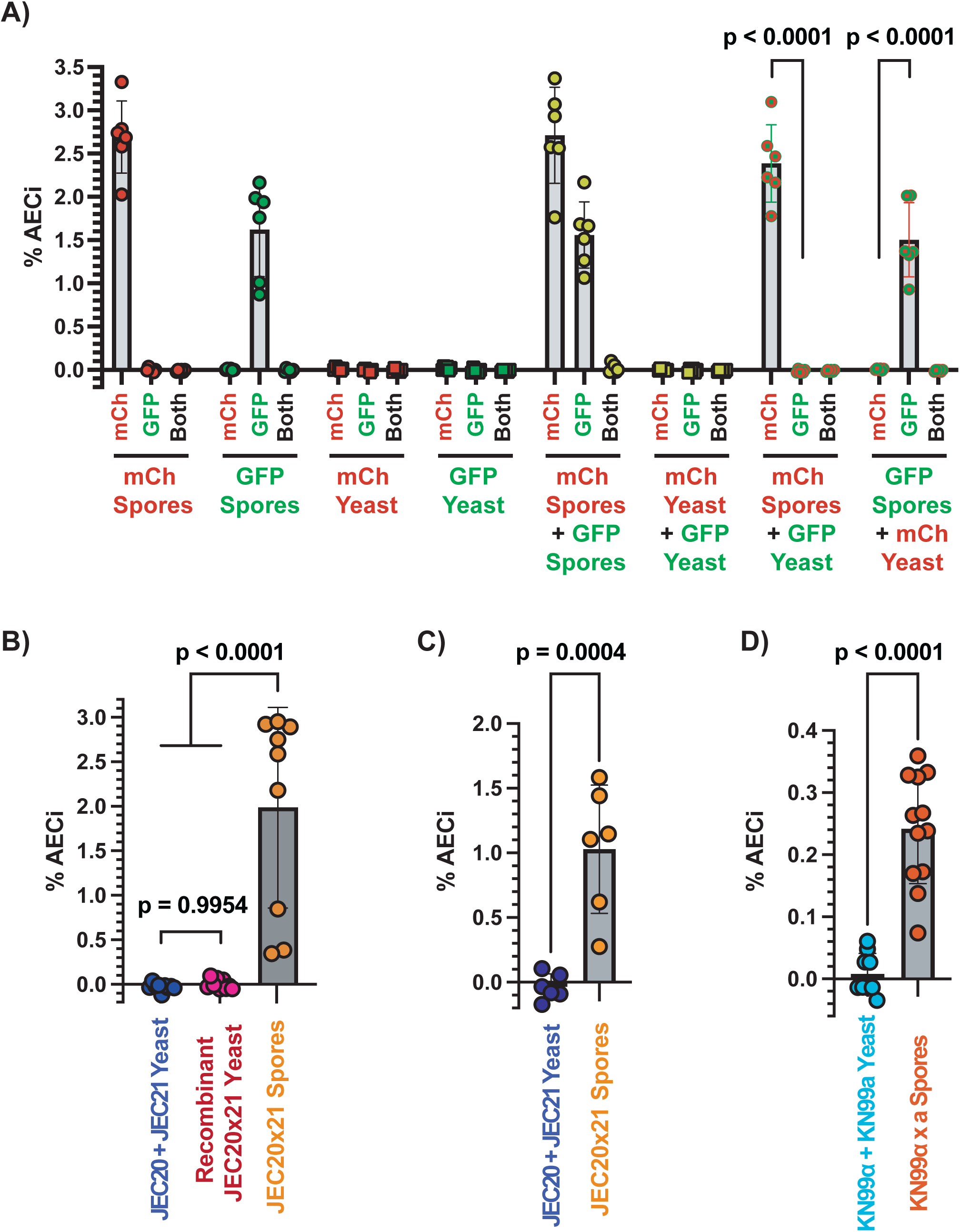
Additional imaging flow cytometry controls validating spore-specific invasion of airway epithelial cells (AECs) by *Cryptococcus*. **A)** Percent of AECs (A549s) with internalization events (AEC_i_) 6 hours post infection (hpi) quantified for infections with either mCh tagged JEC20×21 spores, GFP tagged JEC20×21 spores, mCh tagged JEC20+21 yeast, GFP tagged JEC20+21 yeast, or combinations of these cells. p-values from one-way ANOVA **B)** Percent AEC_i_ 6 hpi for infections of A549s with JEC20×21 spores, JEC20+21 yeast or JEC20×21 germinated spores (i.e. yeast). p-values from one-way ANOVA **C)** Percent AEC_i_ 6 hpi for infections of primary AECs 6 hpi with either JEC20×21 spores or JEC20+21 yeast. p-values from unpaired Student’s T-test **D)** Percent AEC_i_ 6 hpi for infections of A549s with KN99“a”xα spores or KN99“a”+α yeast. p-values from unpaired Student’s T-test

**Supplemental Figure 2.**
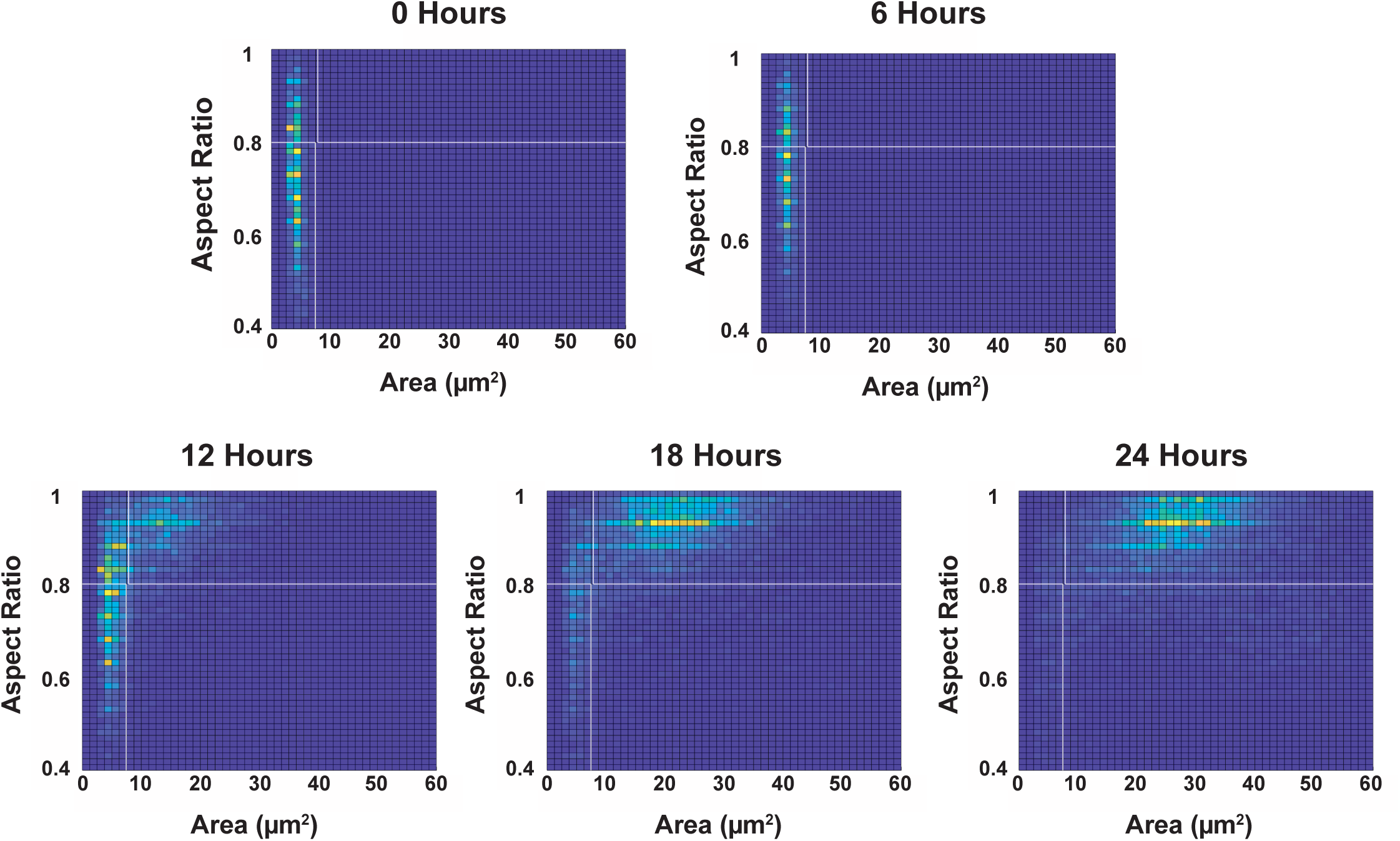
Germination of spores in tissue culture conditions. Two-dimensional histogram of spores (JEC20mChxJEC21mCh) germinated in supplemented DMEM, at 37°C, 5%CO_2_ for 24 hours. Spores are small and oval (Area <7.44 µm^2^, Aspect Ratio <0.8) and present in the lower left hand quadrant, while yeast are large and round (Area >7.84 µm^2^, Aspect Ratio >0.8) and are present in the upper right hand quadrant. Population displays a ‘Bimodal’ phenotype, with the population fully germinated by 24 hours.

**Supplemental Figure 3.**
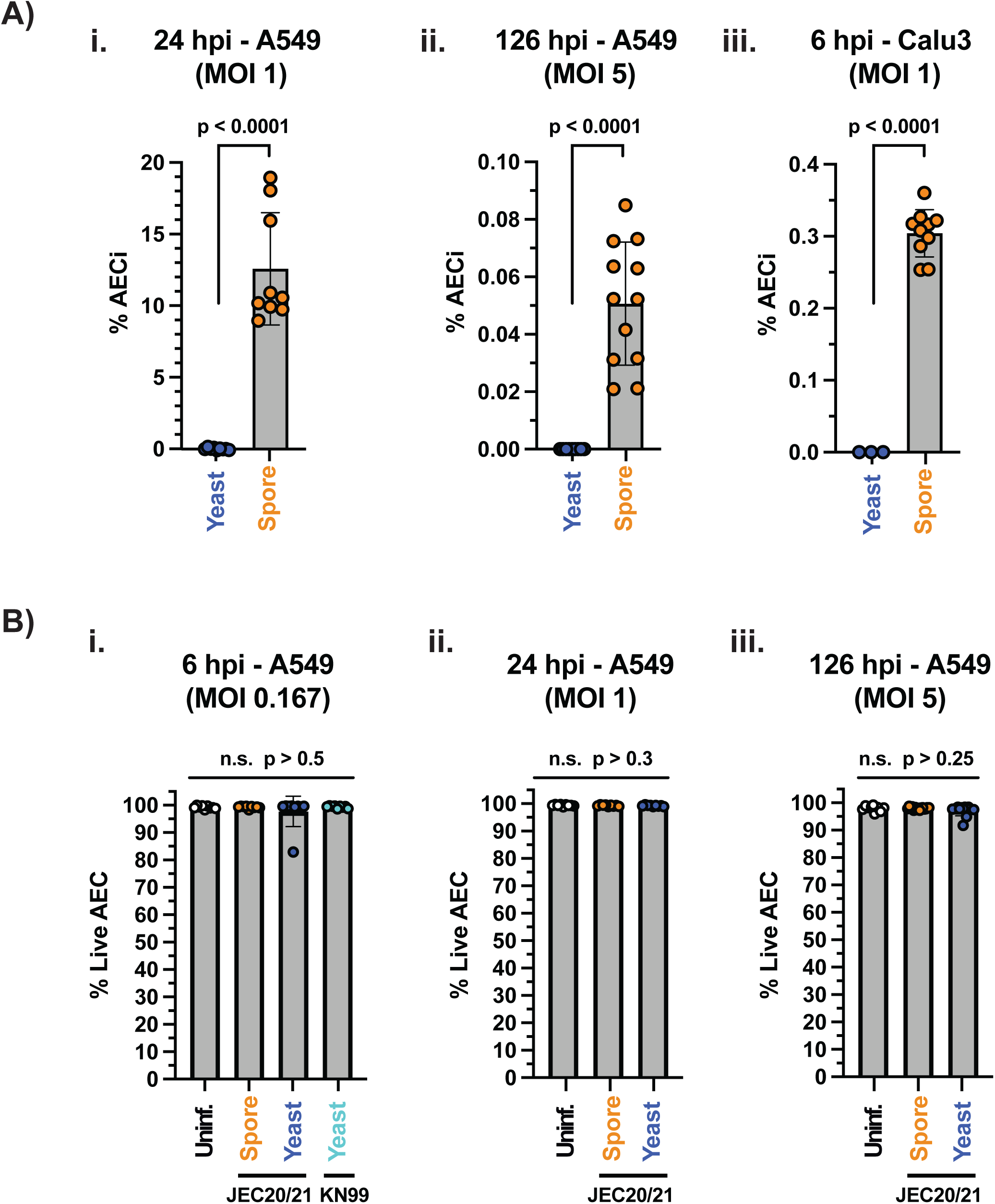
Relative *Cryptococcus* internalisation and host cell damage. **A)** Percent of AECs with intracellular *Cryptococcus* (%AEC_i_) for spore and yeast infected i) A549s for 24 hours - MOI of 1, ii) A549s for 126 hours - MOI of 5, and iii) Calu-3s for 6 hours - MOI of 1. All statistics derived from unpaired Student’s T-test **B)** Percentage of live AECs as determined by lack of necrosis (TO-PRO-3) or apoptosis (Annexin-FITC) for A549s infected for i) 6 hours with an MOI of 0.167, ii) 24 hours with an MOI of 1, or 126 hours with an MOI of 5. All statistics derived from one-way ANOVA across samples.

**Supplemental Figure 4.**
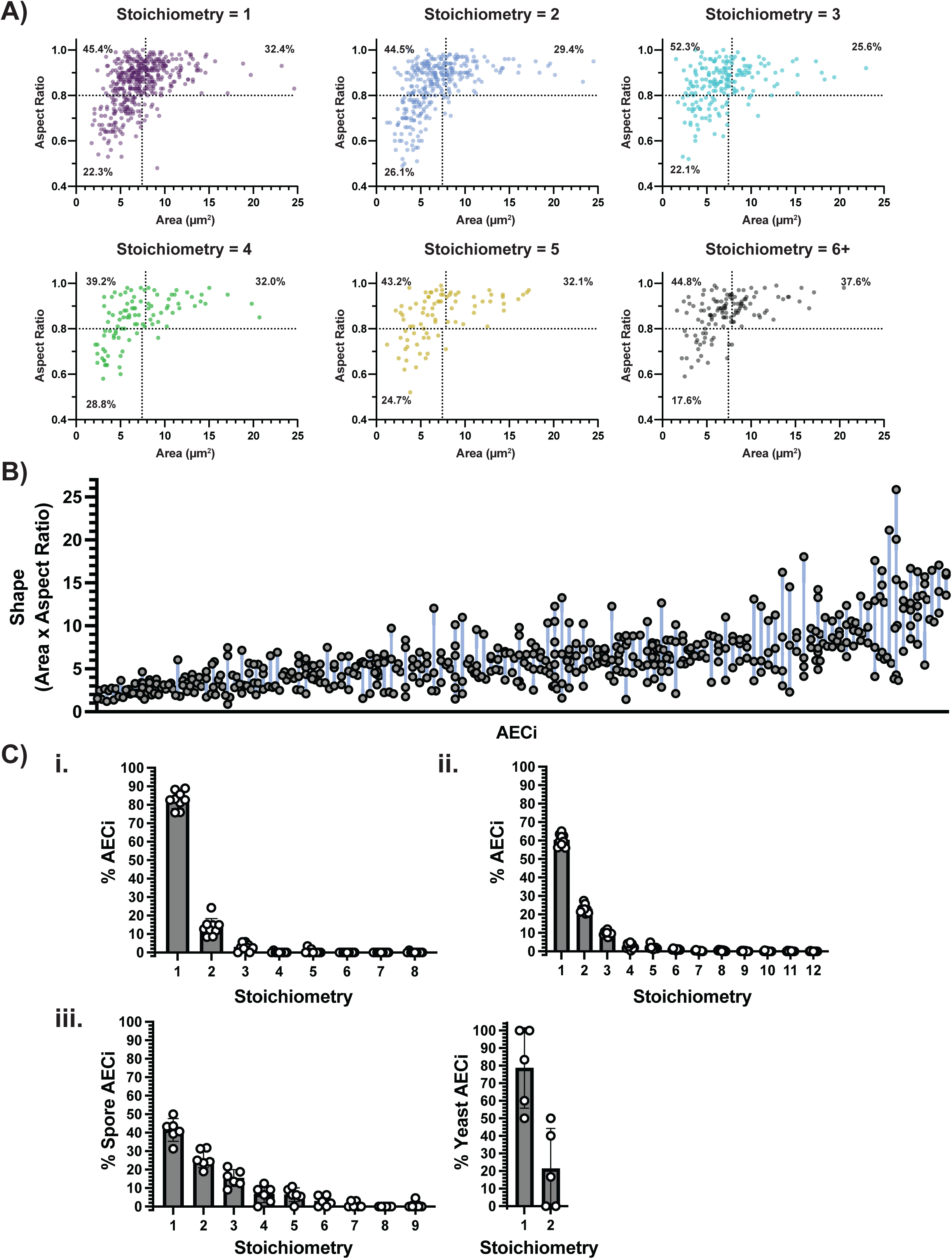
Intracellular *Cryptococcus* stoichiometry. **A)** Germination profiles of intracellular *Cryptococcus* 24 hours post infection separated by number of fungal cells per AEC (stoichiometry) demonstrating that germination kinetics remain constant across all intracellular stoichiometries. **B)** Correlation of spore shape for *Cryptococcus* residing within the same AECs (Stoichiometry = 3+) demonstrate a link between host AEC and germination kinetics p < 0.0001 two-way ANOVA. **C)** Stoichiometry, as percent of population, for i) 6 hour post infection of A549 with spores (MOI 0.167), ii) 24 hours post infection with spores of A549s (MOI 1), and iii) AECs isolated from mice intranasally infected with either spores or yeast.

**Supplemental Figure 5.**
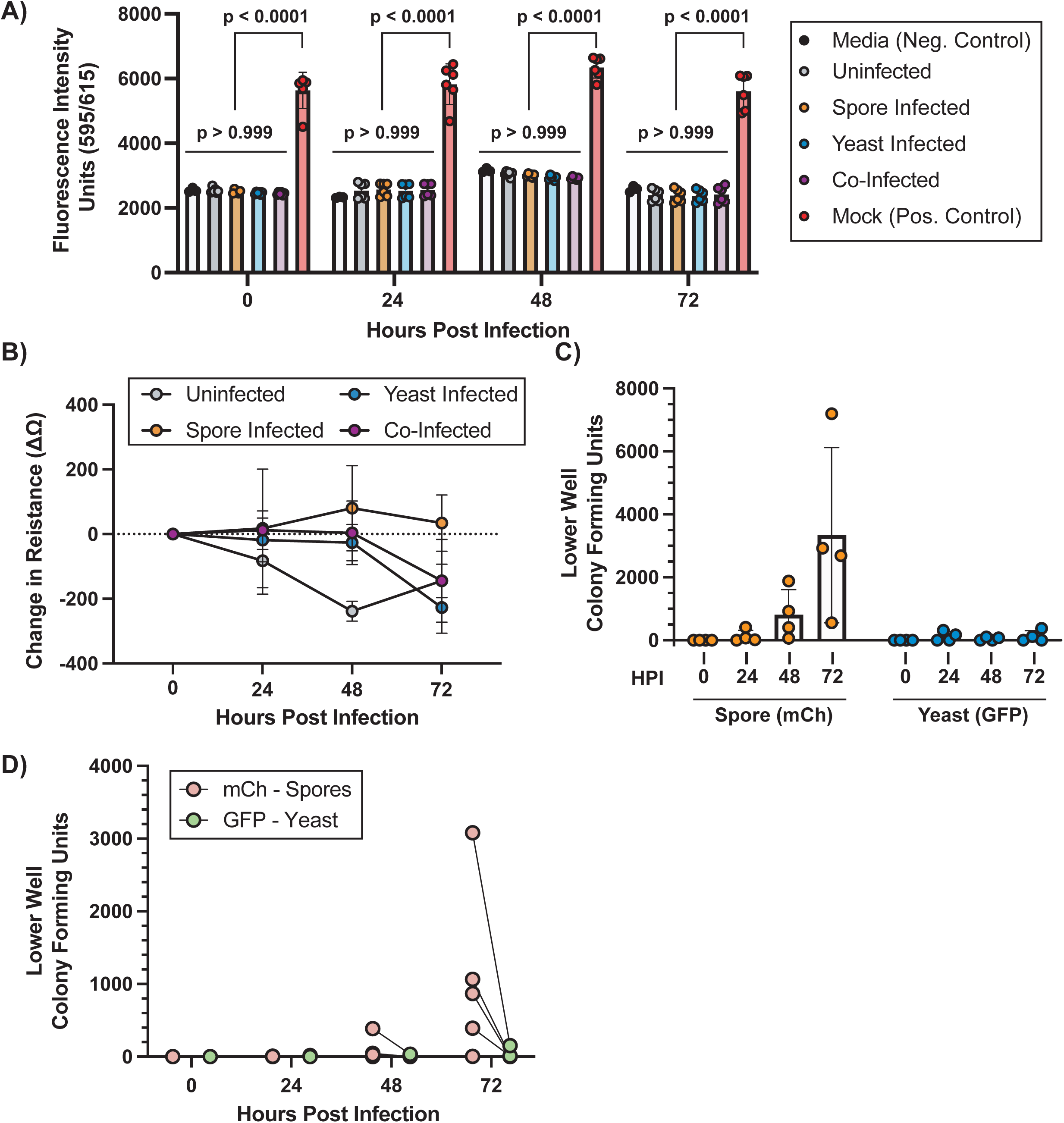
Additional controls for preferential crossing of spores across AEC (Calu-3) barrier. **A)** Fluorescence measurements (595/615nm) of lower well, monitoring the crossing of 70kDa Dextran-Texas Red to evaluate barrier integrity of transwells for uninfected, spore-infected (JEC20×21-mCh), yeast infected (JEC20+21-GFP) and co-infected (JEC20×21-mCh & JEC20+21-GFP) wells. Media (DMEMF12) used as a negative control and mock wells lacking AECs to provide impermeable barriers used as a positive control. All statistics derived from one-way ANOVA **B)** Difference in resistance (Ω) measurements as compared to 0 hpi for uninfected, spore-infected (JEC20×21-mCh), yeast infected (JEC20+21-GFP) and co-infected (JEC20×21-mCh & JEC20+21-GFP) wells as an additional measure of barrier integrity. **C)** Lower well colony forming units at 0, 24, 48 and 72 hours in spore (JEC20×21mCh) and yeast (JEC20+21-GFP) infected tranwells indicating crossing of AEC (Calu-3) barrier, addition of 70kDa Dextan-Texas Red to upper well for barrier integrity monitoring. **D)** Lower well colony forming units at 0, 24, 48 and 72 hours in transwells infected with both spore (JEC20×21mCh) and yeast (JEC20+21-GFP) indicating crossing of AEC (Calu-3) barrier. Spore (mCherry) and yeast (GFP) crossing in the same well are joined by a line. Addition of 70kDa Dextan-Texas Red to upper well for barrier integrity monitoring.

## Citations

[1] Bongomin, F., Gago, S., Oladele, R. O., & Denning, D. W. (2017). Global and Multi-National Prevalence of Fungal Diseases-Estimate Precision. Journal of Fungi (Basel, Switzerland), 3(4), 57. 10.3390/jof3040057

[2] Brown, G. D., Denning, D. W., Gow, N. A., Levitz, S. M., Netea, M. G., & White, T. C. (2012). Hidden killers: human fungal infections. Science Translational Medicine, 4(165), 165rv13. 10.1126/scitranslmed.3004404

[3] World Health Organization. WHO fungal priority pathogens list to guide research, development and public health action. WHO https://www.who.int/publications/i/item/9789240060241 (2022).

[4] Rajasingham, R., Smith, R. M., Park, B. J., Jarvis, J. N., Govender, N. P., Chiller, T. M., Denning, D. W., Loyse, A., & Boulware, D. R. (2017). Global burden of disease of HIV-associated cryptococcal meningitis: an updated analysis. The Lancet. Infectious Diseases, 17(8), 873–881. 10.1016/S1473-3099(17)30243-8

[5] Dromer, F., Casadevall, A., Perfect, J. and Sorrell, T. (2010). *Cryptococcus neoformans*: Latency and Disease. In Cryptococcus (eds J. Heitman, T.R. Kozel, K.J. Kwon-Chung, J.R. Perfect and A. Casadevall). 10.1128/9781555816858.ch31

[6] Alanio A. (2020). Dormancy in *Cryptococcus neoformans*: 60 years of accumulating evidence. The Journal of Clinical Investigation, 130(7), 3353–3360. 10.1172/JCI136223

[7] World Health Organization. Guidelines for diagnosing, preventing and managing cryptococcal disease among adults, adolescents and children living with HIV. WHO https://www.who.int/publications/i/item/9789240052178 (2022).

[8] Ortiz, S. C. and Hull, C.M. 2023. Biogenesis, germination, and pathogenesis of *Cryptococcus* spores. Microbiology and Molecular Biology Reviews. Under Review 2023

[9] Botts, M. R., Giles, S. S., Gates, M. A., Kozel, T. R., & Hull, C. M. (2009). Isolation and characterization of *Cryptococcus neoformans* spores reveal a critical role for capsule biosynthesis genes in spore biogenesis. Eukaryotic Cell, 8(4), 595–605.

[10] Giles, S. S., Dagenais, T. R., Botts, M. R., Keller, N. P., & Hull, C. M. (2009). Elucidating the pathogenesis of spores from the human fungal pathogen *Cryptococcus neoformans*. Infection and Immunity, 77(8), 3491–3500.

[11] Ortiz, S.C., McKeon, M.C., Botts, M. R., Gage, H., Frerichs, A. B & Hull, C. M. (2003) Spores of the fungal pathogen *Cryptococcus* exhibit cell type-specific carbon source utilization during germination. *Preprint at bioRxiv*, 10.1101/2023.10.01.560341

[12] Davis, J. M., Huang, M., Botts, M. R., Hull, C. M., & Huttenlocher, A. (2016). A zebrafish model of cryptococcal infection reveals roles for macrophages, endothelial cells, and neutrophils in the establishment and control of sustained fungemia. Infection and Immunity, 84(10), 3047–3062.

[13] Walsh, N. M., Botts, M. R., McDermott, A. J., Ortiz, S. C., Wüthrich, M., Klein, B., & Hull, C. M. (2019). Infectious particle identity determines dissemination and disease outcome for the inhaled human fungal pathogen *Cryptococcus*. PLoS Pathogens, 15(6), e1007777.

[14] Crapo, J. D., Barry, B. E., Gehr, P., Bachofen, M., & Weibel, E. R. (1982). Cell number and cell characteristics of the normal human lung. The American Review of Respiratory Disease, 125(6), 740–745. 10.1164/arrd.1982.125.6.740

[15] Bertuzzi, M., Hayes, G. E., Icheoku, U. J., van Rhijn, N., Denning, D. W., Osherov, N., & Bignell, E. M. (2018). Anti-*Aspergillus* activities of the respiratory epithelium in health and disease. Journal of Fungi, 4(1), 8. 10.3390/jof4010008

[16] Bertuzzi, M., Hayes, G. E., & Bignell, E. M. (2019). Microbial uptake by the respiratory epithelium: outcomes for host and pathogen. FEMS Microbiology Reviews, 43(2), 145–161. 10.1093/femsre/fuy045

[17] Bertuzzi, M., Howell, G.J., Thomson, D.D., Fortune-Grant, R., Möslinger, A., Dancer, P., Van Rhijn, N., Du, X., Codling, A., Sash, R., Demirbag, M., Bignell, E.M. (2022) Epithelial uptake of *Aspergillus fumigatus* drives efficient fungal clearance *in vivo* and is aberrant in Chronic Obstructive Pulmonary Disease (COPD). Preprint at bioRxiv, 10.1101/2022.02.01.478664

[18] Merkel, G. J., & Scofield, B. A. (1997). The in vitro interaction of *Cryptococcus neoformans* with human lung epithelial cells. FEMS Immunology and Medical Microbiology, 19(3), 203–213. 10.1111/j.1574-695X.1997.tb01089.x

[19] Barbosa, F. M., Fonseca, F. L., Holandino, C., Alviano, C. S., Nimrichter, L., & Rodrigues, M. L. (2006). Glucuronoxylomannan-mediated interaction of *Cryptococcus neoformans* with human alveolar cells results in fungal internalization and host cell damage. Microbes and Infection, 8(2), 493–502. 10.1016/j.micinf.2005.07.027

[20] Choo, K. K., Chong, P. P., Ho, A. S., & Yong, P. V. (2015). The role of host microfilaments and microtubules during opsonin-independent interactions of *Cryptococcus neoformans* with mammalian lung cells. European Journal of Clinical Microbiology & Infectious Diseases, 34(12), 2421–2427. 10.1007/s10096-015-2497-4

[21] Taylor-Smith L. M. (2017). *Cryptococcus*-epithelial interactions. Journal of Fungi (Basel, Switzerland), 3(4), 53. 10.3390/jof3040053

[22] Walsh, N. M., Wuthrich, M., Wang, H., Klein, B., & Hull, C. M. (2017). Characterization of C-type lectins reveals an unexpectedly limited interaction between *Cryptococcus neoformans* spores and Dectin-1. PloS One, 12(3), e0173866. 10.1371/journal.pone.0173866

[23] Bertuzzi, M., & Howell, G. J. (2021). Single-cell analysis of fungal uptake in cultured airway epithelial cells using differential fluorescent staining and imaging flow cytometry. Methods in Molecular Biology (Clifton, N.J.), 2260, 83–109. 10.1007/978-1-0716-1182-1_6

[24] Ortiz, S. C., Huang, M., & Hull, C. M. (2021). Discovery of fungus-specific targets and inhibitors using chemical phenotyping of pathogenic spore germination. mBio, 12(4), e0167221. 10.1128/mBio.01672-21

[25] Barkal, L. J., Walsh, N. M., Botts, M. R., Beebe, D. J., & Hull, C. M. (2016). Leveraging a high resolution microfluidic assay reveals insights into pathogenic fungal spore germination. Integrative Biology, 8(5), 603–615. 10.1039/c6ib00012f

[26] Okagaki, L. H., Strain, A. K., Nielsen, J. N., Charlier, C., Baltes, N. J., Chrétien, F., Heitman, J., Dromer, F., & Nielsen, K. (2010). *Cryptococca*l cell morphology affects host cell interactions and pathogenicity. PLoS Pathogens, 6(6), e1000953. 10.1371/journal.ppat.1000953

[27] Ma, H., Croudace, J. E., Lammas, D. A., & May, R. C. (2006). Expulsion of live pathogenic yeast by macrophages. Current Biology : CB, 16(21), 2156–2160. 10.1016/j.cub.2006.09.032

[28] Alvarez, M., & Casadevall, A. (2006). Phagosome extrusion and host-cell survival after *Cryptococcus neoformans* phagocytosis by macrophages. Current Biology, 16(21), 2161–2165. 10.1016/j.cub.2006.09.061

[29] Tucker, S. C., & Casadevall, A. (2002). Replication of *Cryptococcus neoformans* in macrophages is accompanied by phagosomal permeabilization and accumulation of vesicles containing polysaccharide in the cytoplasm. Proceedings of the National Academy of Sciences of the United States of America, 99(5), 3165–3170. 10.1073/pnas.052702799

[30] Lin, X., Nielsen, K., Patel, S., & Heitman, J. (2008). Impact of mating type, serotype, and ploidy on the virulence of *Cryptococcus neoformans*. Infection and Immunity, 76(7), 2923–2938. 10.1128/IAI.00168-08

[31] Sutherland, T. C., Quattroni, P., Exley, R. M., & Tang, C. M. (2010). Transcellular passage of *Neisseria meningitidis* across a polarized respiratory epithelium. Infection and immunity, 78(9), 3832–3847. 10.1128/IAI.01377-09

[32] Francis, V. I., Liddle, C., Camacho, E., Kulkarni, M., Junior S.R.S., Harvey, J.A., Ballou, E.R., Thomson, D.D., Brown, G. D., Hardwick J.M., Casadevall, A., Witton, J., & Coelho, C. (2024). *Cryptococcus neoformans* rapidly invades the murine brain by sequential breaching of airway and endothelial tissues barriers, followed by engulfment by microglia. mBio, 15(4), e0307823. 10.1128/mbio.03078-23

[33] Bermudez, L. E., Sangari, F. J., Kolonoski, P., Petrofsky, M., & Goodman, J. (2002). The efficiency of the translocation of *Mycobacterium tuberculosis* across a bilayer of epithelial and endothelial cells as a model of the alveolar wall is a consequence of transport within mononuclear phagocytes and invasion of alveolar epithelial cells. Infection and Immunity, 70(1), 140–146. 10.1128/IAI.70.1.140-146.2002

[34] Garcia-Hermoso, D., Janbon, G., & Dromer, F. (1999). Epidemiological evidence for dormant *Cryptococcus neoformans* infection. Journal of Clinical Microbiology, 37(10), 3204–3209. 10.1128/JCM.37.10.3204-3209.1999

[35] Ristow, L. C., & Davis, J. M. (2021). The granuloma in cryptococcal disease. PLoS Pathogens, 17(3), e1009342. 10.1371/journal.ppat.1009342

[36] Mayito, J., Andia, I., Belay, M., Jolliffe, D. A., Kateete, D. P., Reece, S. T., & Martineau, A. R. (2019). Anatomic and cellular niches for *Mycobacterium tuberculosis* in latent tuberculosis infection. The Journal of Infectious Siseases, 219(5), 685–694. 10.1093/infdis/jiy579

[37] Hernández-Pando, R., Jeyanathan, M., Mengistu, G., Aguilar, D., Orozco, H., Harboe, M., Rook, G. A., & Bjune, G. (2000). Persistence of DNA from *Mycobacterium tuberculosis* in superficially normal lung tissue during latent infection. Lancet (London, England), 356(9248), 2133–2138. 10.1016/s0140-6736(00)03493-0

[38] Alanio, A., Vernel-Pauillac, F., Sturny-Leclère, A., & Dromer, F. (2015). *Cryptococcus neoformans* host adaptation: toward biological evidence of dormancy. mBio, 6(2), e02580–14. 10.1128/mBio.02580-14

[39] Hommel, B., Sturny-Leclère, A., Volant, S., Veluppillai, N., Duchateau, M., Yu, C. H., Hourdel, V., Varet, H., Matondo, M., Perfect, J. R., Casadevall, A., Dromer, F., & Alanio, A. (2019). *Cryptococcus neoformans* resists to drastic conditions by switching to viable but non-culturable cell phenotype. PLoS pathogens, 15(7), e1007945. 10.1371/journal.ppat.1007945

[40] Upadhya, R., Lam, W. C., Maybruck, B. T., Donlin, M. J., Chang, A. L., Kayode, S., Ormerod, K. L., Fraser, J. A., Doering, T. L., & Lodge, J. K. (2017). A fluorogenic *C. neoformans* reporter strain with a robust expression of m-cherry expressed from a safe haven site in the genome. Fungal Genetics and Biology, 108, 13–25. 10.1016/j.fgb.2017.08.008

[41] Sherman F, Fink GR, Hicks JB. (1987). Laboratory course manual for methods in yeast genetics. Cold Spring Harbor Laboratory, Cold Spring Harbor, NY.

